# Accuracy and Power Analysis of Social Networks Built From Count Data

**DOI:** 10.1101/2021.05.07.443094

**Authors:** Jordan D. A. Hart, Daniel W. Franks, Lauren J. N. Brent, Michael N. Weiss

## Abstract

1. Power analysis is used to estimate the probability of correctly rejecting a null hypothesis for a given statistical model and dataset. Conventional power analyses assume complete information, but the stochastic nature of behavioural sampling can mean that true and estimated networks are poorly correlated. Power analyses do not currently take the effect of sampling into account. This could lead to inaccurate estimates of statistical power, potentially yielding misleading results.
2. Here we develop a method for computing *network correlation*: the correlation between an estimated social network and its true network, using a Gamma-Poisson model of social event rates for networks constructed from count data. We use simulations to assess how the level of network correlation affects the power of nodal regression analyses. We also develop a generic method of power analysis applicable to any statistical test, based on the concept of diminishing returns.
3. We demonstrate that our network correlation estimator is both accurate and moderately robust to its assumptions being broken. We show that social differentiation, mean social event rate, and the harmonic mean of sampling times positively impacts the strength of network correlation. We also show that the required level of network correlation to achieve a given power level depends on many factors, but that 0.80 network correlation usually corresponds to around 0.80 power for nodal regression in ideal circumstances.
4. We provide guidelines for using our network correlation estimator to verify the accuracy of networks built from count data, and to conduct power analysis. This can be used prior to data collection, in post hoc analyses, or even for subsetting networks in dynamic network analysis. The network correlation estimator and custom power analysis methods have been made available as an R package.

## Introduction

Understanding the form and function of social systems is a core aim of behavioural ecology, and social network analysis has become a central tool for exploring these topics (Croft et al., 2008). However, social network analysis comes with challenges unique to the field, ranging from issues of missing or censored data, through to potential problems of non-independence in network measures (James et al., 2009; Farine and Whitehead, 2015). Significant advances have been made in many of these areas over the last two decades, but the problem of conducting power Whitehead analysis has generally received little attention, with the exception of (2008). Power analysis is used to determine the amount of data required to reject a null hypothesis assuming a given effect size, and is used to determine how a study should be conducted prior to collecting data, or in post-hoc analysis to determine if a study was sufficiently powered to reject a null hypothesis (Cohen, 1992, 2013). Animal social network studies rarely use power analysis prior to data collection because there are often many unknown variables, and sample sizes are often fixed. However, they often do employ post hoc power analysis to indicate the reliability of results (Martin et al., 1993; Stadtfeld et al., 2020). In this study, we build on previous work to develop a method for estimating the accuracy of social networks, and show how this can be used to conduct power analysis on social networks constructed from count data.

Networks are typically built from weighted edges between nodes that encode the strengths of relationships between individuals (Krause et al., 2015). Edge weight can be quantified in a number of different ways depending on the nature of the social system and the data available. The most common method for calculating network edges is to compute the simple ratio index, which normalises a measure of sociality against sampling time. Sociality is usually quantified by recording *social events* such as spatial associations or social interactions. There are three ways social events are often recorded, which we will refer to as binary, count, and duration. Binary data records the presence or absence of a social event in each of a series of fixed sampling intervals. Social events can only be marked as present or absent in each interval. Count data records the number of social events observed over the amount of time spent sampling. Finally, duration data records the amount of time spent engaged in a social event over the amount of time spent sampling (Croft et al., 2008). In networks built from binary data, the simple ratio index is equivalent to the probability of observing a social event in a given sampling period. Whereas in networks built from count data, the simple ratio index is the rate of social events per unit of time.

Whitehead (2008) introduced a method for estimating the accuracy of animal social networks constructed from binary data by estimating the correlation of the observed event probabilities with the underlying probabilities. This method has often been used as a post hoc measure to verify the robustness of animal social networks before further analyses (Findlay et al., 2016; Ellis et al., 2017). However, this method is designed specifically for binary data and not count or duration data. In this work we develop a variation of this method for social networks constructed from count data.

The correlation between true and estimated networks, which we will refer to as *network correlation*, is a useful intuitive measure of the accuracy of a network, but even more importantly it is directly related to the power of statistical analyses. Whitehead (2008) made a series of recommendations based on the social preference test developed by Bejder et al. (1998), suggesting that a good general guide for maximal power is when the product of the squared *social differentiation* and mean number of observations per individual is greater than 5 (in their notation: *S*^2^ × *H >* 5). In this context, social differentiation is defined as the coefficient of variation of edge weights. This recommendation is based only on the social preference test, and it isn’t known how well this generalises to other statistical tests commonly used in social network analysis (Bejder et al., 1998). In particular, one of the most common methods for testing hypotheses in social network analysis is nodal regression (Croft et al., 2011). Nodal regression uses a node-level social network metric such as node strength, eigenvector centrality, or closeness to quantify an individual’s position in their social structure, and relates this social position to quantifiable traits such as age or sex (Farine and Whitehead, 2015). The relationship between network metric and trait is usually analysed using regression (Weiss et al., 2021b). The statistical power of a conventional regression depends on sample size, effect size, and significance level. In a nodal regression, sample size is the number of individuals, the true effect size is fixed and unknown, and the significance level is set by convention, usually to 0.05 (Cohen, 1994). However, the true effect size is the effect size if the nodal regression was run on the true, unknown network. The estimated network will not perfectly match the true network, meaning that in turn the estimated effect size will deviate from the true effect size. Applying conventional power analysis to nodal regression could therefore lead to under- or over-estimates of power, potentially yielding misleading results. In general, greater noise in predictor variables reduces the magnitude of coefficient estimates, increasing the chance of significant effects being missed when they are in fact present (Frost and Thompson, 2000)

In this study we extend the method developed by Whitehead (2008) for estimating network correlation from networks constructed from association data to networks constructed from count data. Count data are usually collected by recording instances of pairs of individuals engaged in a social event over many sampling periods using either scan or focal sampling (Martin et al., 1993). The result is an integer count of social events *X*_*i j*_ between each pair of individuals *i* and *j*, and a positive real-valued sampling time for each pair *d*_*i j*_. The sampling time is the amount of time where at least one of the pair was visible to the observer, and therefore a social event between *i* and *j* could have taken place. The rate of social events is then computed by dividing the social event count by the sampling time, giving the social event rate:

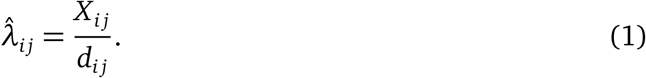

As with any empirical measure, sampling has the potential to influence the estimated social event rate significantly (Franks et al., 2010). Consider the case when *X*_*i j*_ is 10 events and *d*_*i j*_ is 5 minutes, if event counts are Poisson-distributed, the estimate will be 2 social events per minute, but the 95% confidence interval will be (0.9, 3.7), with a range of 2.6. Compare this to the case where *X*_*i j*_ and *d*_*i j*_ are 100 events and 50 minutes respectively; the point estimate is the same, but the 95% confidence interval is (1.6, 2.4), with a range of 0.8, less than one third of the range of the previous case, but for around 10 times the sampling time. In this example we used minutes as the unit of time, but in general the units of time do not matter as long as they are consistent. This shows that we should have less confidence in the estimate of the first case compared to the second case, and demonstrates how the estimated network could deviate significantly from the true network because of low sampling time. The magnitude of this deviation will impact the reliability of an analysis, and ideally should be taken into account when conducting social network analyses. This problem has been recognised in several previous studies (Whitehead, 2008; Lusseau et al., 2008; Farine and Strandburg-Peshkin, 2015; Davis et al., 2018). In particular, Whitehead (2008) developed a method based on the binomial distribution for estimating network correlation for networks constructed from binary data.

We demonstrate that our method for count data provides accurate estimates of network correlation in realistic scenarios, and we use simulations to suggest guidelines for the minimum level of sampling required for nodal regression depending on the desired level of statistical power. We also develop a generic alternative approach for guiding data collection based on the principle of diminishing returns that can be used for any type of statistical analysis, including but not limited to nodal regression. We contrast our guidelines to those from Whitehead (2008) to show that the amount of sampling required depends on the type of data, and that data-specific guidelines may be highly useful when designing and conducting social network analysis. We have made the methods available as an R package: *pwrCGP*, which is available at www.github.com/JHart96/pwrCGP.

## Methods

By modelling event counts as being distributed according to a Gamma-Poisson process, we are able to analytically derive an equation for network correlation. We verify our network correlation equation using simulations which either follow the assumptions of the model, or break them to varying degrees, to test the robustness of the method. Following this, we use simulations to determine the level of network correlation required to obtain a desired level of power when performing nodal regression on networks built from count data.

### Gamma-Poisson model of social event rates

As sampling time increases, we expect that the estimated social event rate 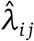 will get closer to the true social event rate *λ*_*i j*_. However for lower sampling times there may be a considerable error between the estimated and true event rate. This error can be modelled by treating the event counts *X*_*i j*_ as draws from a Poisson distribution. Since the underlying true event rates *λ*_*i j*_ are unknown, we assume the true event rates of the dyads are drawn from a gamma distribution: *λ*_*i j*_ ∼ Gamma(*α, β*). A Poisson-distributed random variable with rates drawn from the gamma distribution is equivalent to a random variable following the negative binomial distribution, therefore the number of observed events *X*_*i j*_ is given by:

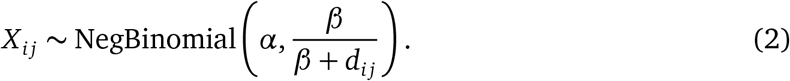

Using this, we can estimate the variance of both the true and estimated event rates, which allows us to estimate the Pearson’s correlation coefficient, *ρ*, between them:

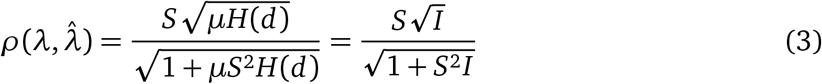

where 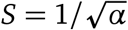 is the social differentiation, *µ* = *α/β* is the dyadic mean social event rate, *H*(*d*) is the harmonic mean of the sampling times *d*_*i j*_, and *I* = *µH*(*d*) reflects the sampling effort (see supplementary materials for full derivation). The harmonic mean *H*(*d*) of the *m* dyads is defined as 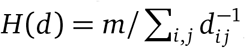, and is equal to the arithmetic mean only in the case where all *d*_*i j*_ are equal. Note that when sampling time *d*_*i j*_ is even across all dyads, sampling effort *I* is the number of social events observed per dyad. When sampling time *d*_*i j*_ is uneven, sampling effort *I* will be lower, and more sampling time will be required to reach the same sampling effort as the equivalent network with evenly-sampled dyads. The network correlation is computed only over dyads that have non-zero sampling times.

The parameters *α, β* of the underlying gamma distribution can be estimated numerically using maximum likelihood. We use point estimates from maximum likelihood in this study to reduce computation time, and to avoid model fitting problems. We have also included a version in the code that uses quadratic approximation to estimate confidence intervals of network correlation.

### Simulations 1: Verification of the Gamma-Poisson model

To confirm that Equation 3 is appropriate for event rate data, and to determine how robust it is to the Gamma-Poisson assumptions being broken, we ran simulations under three different scenarios: 1) following the assumptions of the model, 2) introducing community structure to the network, and 3) having zero-inflated edge weights (see supplementary material for more details). The parameter space of these scenarios was explored using a random search for *S* ∈ (0, 2] and *µ* ∈ (0, 10]. For scenario 2, an additional constraint was applied such that event rates are stronger between members of the same group than between members of different groups. The simulations proceeded as follows:

1. Data were generated according to one of the scenarios.
2. The true correlation between *λ* and 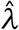 was computed.
3. The parameters *µ* and *S* were estimated using maximum likelihood and used to compute the estimated correlation 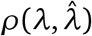 between *λ* and 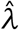.
4. The network correlation was estimated using Equation 3, and the process was repeated 200 times.

The relationship between the true and estimated network correlations was quantified by computing the Pearson correlation coefficient between the estimated network correlations and true network correlations, the mean absolute error, and the standard deviation of the error.

### Simulations 2: Statistical power of nodal regression

We ran simulations to test how statistical power relates to sampling effort in the case where the true effect size is the minimum required to achieve 100% power. This made it possible to see the effect of sampling effort without the effect being hidden behind under- or over-powered tests. To simulate data collection we followed the core assumptions of the model: that dyad-level event rates are drawn from a gamma distribution, and observations are made such that event counts follow a Poisson distribution. Sampling time for each dyad was modelled as the number of sampling periods, which was drawn from a Poisson distribution with mean *D* ∈ [10, 10000]. The dyad-level event rates were converted to an edge list from which the true network was built. Node strength *s*_*i*_ was used as the node-level network metric for these simulations. A linear relationship between network metric and individual trait was created by assigning traits *t*_*i*_ to each individual *i* according to a linear equation *t*_*i*_ = *a* + *bs*_*i*_ + *ε*_*i*_, where *a* is the equivalent of an intercept term, *b* encodes the relationship between metric and trait, and *ε*_*i*_ is a normally-distributed noise term: *ε*_*i*_ ∼ *N* (0, 1). The value of the effect *b* was set depending on the number of nodes *n*. This was determined using a preliminary simulation in such a way that the value of *b* for a given number of individuals was equal to the lowest value for which a power ≥ 99.9% can be achieved.

Networks were estimated using Equation 1 and node strength was estimated from the corresponding estimated networks. To estimate the relationship *b*, a simple linear regression was fitted to the simulated data, and by convention, node-label permutations were used to calculate the p-values (R Core Team, 2013; Croft et al., 2011). The power of the nodal regression was computed for each set of parameters by repeatedly assigning traits *t*_*i*_ according to the true network, fitting a linear model to an estimated network, and computing the p-value of the estimated relationship. The proportion of p-values less than 0.05 gave the power of the test for the current set of parameters. We searched the parameter space using a random search to assess the relationship between network correlation, statistical power, and the number of individuals.

Finally, to provide guidelines on the level of network correlation required in a nodal regression, LOESS curves approximating the relationship between correlation and power for different numbers of individuals were fitted to the simulated data (Cleveland, 1979; R Core Team, 2013). The resulting curves were used to predict the level of network correlation required to achieve 80% power, assuming the underlying relationship has power ≈ 100%.

### Simulations 3: Optimal network correlation estimator for generic tests

The relationship between network correlation (Equation 3) and statistical power is affected by several factors, many of which will depend on the type of analysis being conducted. The relationship between power and network correlation was explored in detail for nodal regression in the previous section, but simulation-based studies like this are limited to the analyses they focus on, and cannot generalise to other methods. However we can expect that as network correlation increases, the power of any statistical tests should also improve (or at least stay the same). Assuming that increases in network correlation positively affect statistical power, and that increases in sampling effort come at a cost to researchers, the problem of finding the optimal sampling effort can be seen as finding the point at which increases in sampling effort lead to diminishing returns.

Diminishing returns describes how the rate of increase of one variable decreases as another variable increases (Shephard and Färe, 1974). In our case the aim is to find the point at which increases in sampling effort lead to diminishing increases in network correlation. This is the same problem as finding the “elbow point” of the relationship between network correlation and sampling effort. There is no guarantee of the power of the analysis at the elbow point, but we do know that additional sampling would provide increasingly small gains in network correlation. Since collecting behavioural data is generally time-consuming, financially expensive, and may even have ethical implications, assuming a cost to increases in sampling effort allows us to use the elbow point as an estimate for the optimal level of network correlation and corresponding amount of sampling (Martin et al., 1993). Though this will not guarantee the power of an analysis, if social differentiation and sampling effort can be estimated well, it will indicate the point at which additional sampling will yield increasingly small increases in network correlation.

The elbow of the curve can be computed numerically (further details are included in the supplementary material), but because *ρ* is asymptotic to 1.0, no true elbow exists. We can get around this problem by introducing a free parameter *ρ*_MAX_ to describe the effective maximum network correlation to be used for computing the elbow. The choice of *ρ*_MAX_ encodes the minimum acceptable trade-off between increases in network correlation and increases in sampling effort, and therefore will affect the estimated elbow point. However, using a value of *ρ*_MAX_ sufficiently high that we would consider a sampled network to be negligibly different to the true network (for biological purposes) represents a meaningful choice since increases in network correlation beyond *ρ*_MAX_ would add no further value for our purposes. We use a value of *ρ*_MAX_ = 0.99 for our analysis, but a brief exploration of the impact of this choice is included in the supplementary material.

To assess the levels of power obtained by using the optimal network correlations, we simulated nodal regression analyses for different levels of social differentiation *S* ∈ (0.0, 0.5] and sampling effort *I* ∈ (0, 500] using a similar setup as the previous section. The power of the analysis for each simulated “true” network was computed, and the proportion of networks with lower than the optimal level of network correlation with power greater than 80% were calculated. This was repeated for the proportion of networks with the same or higher than the optimal level of network correlation. This provides a descriptive measure of the performance of the method for estimating the sampling required to achieve 80% power. This level of power was chosen from convention, but there is no reason that the optimal network correlation estimator should obey this convention as it is based on a different concept to conventional power analysis. We also estimated the level of power that most closely matched the estimator for nodal regression analysis using a grid search.

### Case study: Southern resident killer whale contacts

To demonstrate our method, we applied it to a publicly-available dataset of near-surface physical contact interactions between 22 southern resident killer whales *(Orcinus orca)*. Interactions were observed using an unoccupied aerial vehicle over a total of 11 hours during the summer of 2019 (Weiss et al., 2021a). The mean number of observed interactions per dyad was 3.43, with the average sampling time per dyad being 210 minutes. We used numerical MLE to estimate the confidence intervals of social differentiation *S* and network correlation *ρ*. This is implemented in the *pwrIRGP* R package in the function `net_cor`. Additionally, we conducted a nodal regression power analysis on the data for different levels of true effect size (*r* ∈ {0.1, 0.3, 0.5, 0.7, 0.9}) using our function `pwr_nodereg`. We also applied the diminishing returns sampling effort estimator using the function `pwr_elbow` to estimate the optimal level of network correlation and its corresponding sampling effort.

## Results

### Gamma-Poisson model of social event rates

The analytical equation for estimated network correlation given by Equation 3 was used to produce the plots in Figure 1 of network correlation against social differentiation and sampling effort. This shows that, as expected, network correlation increases towards an asymptote at 1.0 with both social differentiation *S* and sampling effort *I*. Lower numbers of social events (*I* = 1) did not reach a network correlation of 1.0 even for high levels of social differentiation (*S* = 1.0). Higher sampling efforts of *I* = 100 reached close to a network correlation of 1.0 even at low levels of social differentiation, *S* < 0.25. This shows that social differentiation is an important factor in estimating the network correlation of a sampled network, which is in line with the findings of (Whitehead, 2008). The results show that relatively low values of *S* ≈ 0.25 are required to make achieving a high level of network correlation feasible, but that lower values of social differentiation (*S* = 0.05) require sampling effort *I >* 100 to achieve even 50% network correlation.

**Figure 1:**
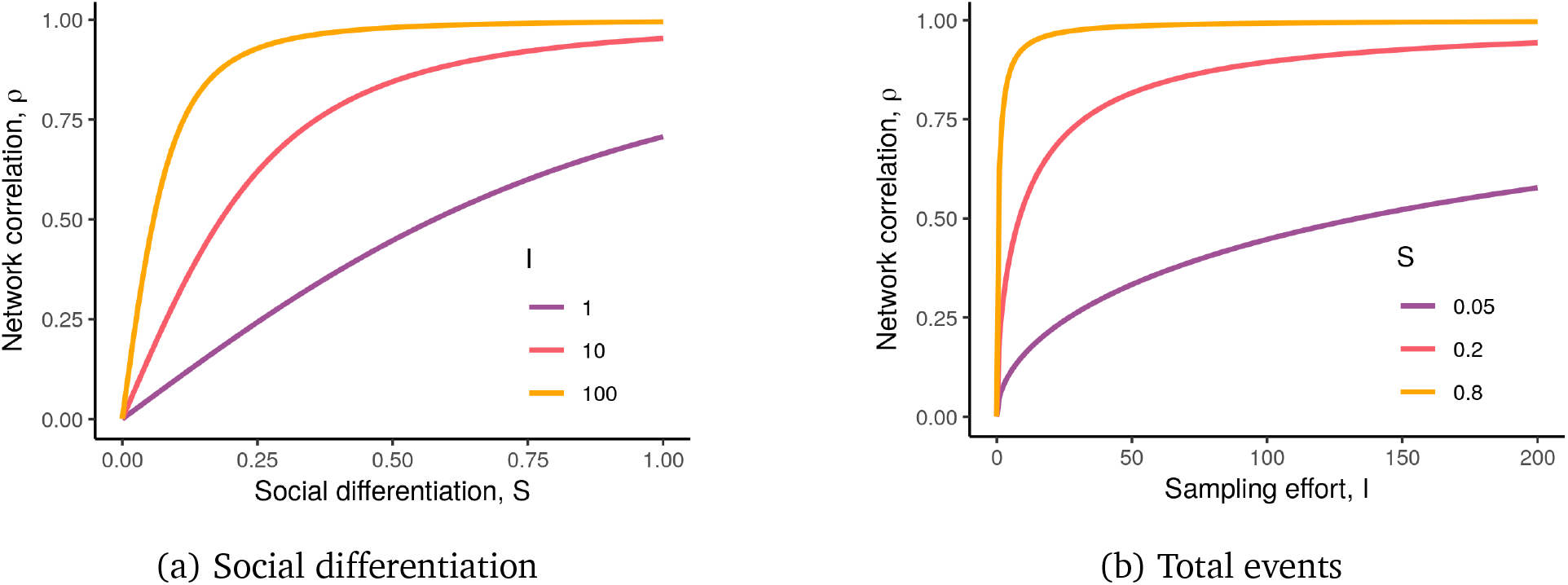
Plots of network correlation against social differentiation *S* and sampling effort *I* = *µH*(*d*) for different levels of sampling and social differentiation respectively using Equation 3. This shows that for relatively low levels of social differentiation around *S* = 0.25, a high level of sampling effort around *I* = 100 is required to achieve a network correlation of *ρ* = 0.90. It also shows that for low social differentiation of *S* = 0.05, even high levels of sampling effort of *I* = 100 will not yield a network correlation of *ρ* = 0.90.

### Simulations 1: Verification of the Gamma-Poisson model

The results of our simulations (see Figure 2) show that true and estimated network correlation match closely across the parameter space. Over the full parameter space, the mean absolute error between the true and estimated network correlations was 5.5%, 0.047%, and 3.5% for scenarios 1, 2, and 3 respectively. The standard deviations of the errors for the three scenarios were 0.0084, 0.0038, and 0.047 respectively, and the correlations between the true and estimated network correlations were 0.99, 0.99, and 0.96 respectively. The relationship between true and estimated network correlation is shown in Figure 2 over a limited part of the parameter space, where the social differentiation was adjusted manually to visualise the full width of the distribution of network correlations. Social differentiation was used to adjust the mean level of network correlation purely for visualisation purposes, and was not used when computing the quantitative statistics.

**Figure 2:**
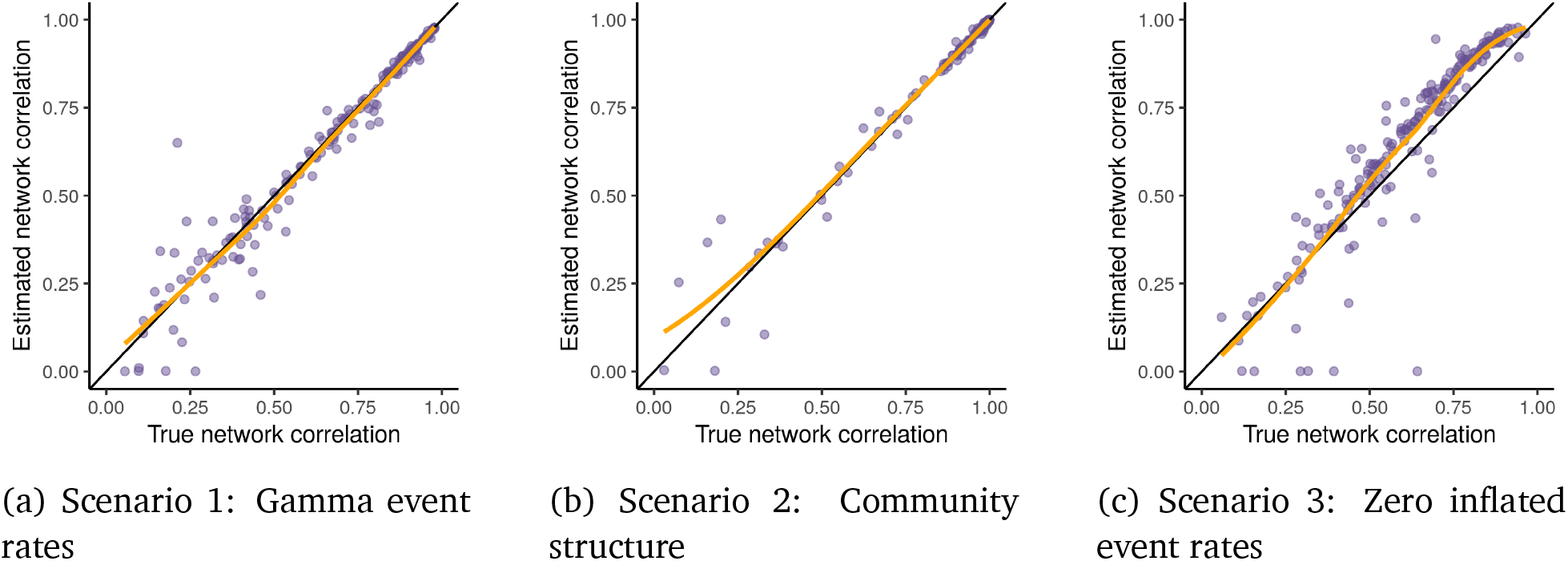
Comparisons of the true network correlations and the network correlations estimated using Equation 3. The black line is diagonal and shows the ideal relationship between the true and estimated network correlations. In these visualisations the three models were run with different maximum social differentiation parameters to ensure the full distribution of network correlations was visible, these were 0.20, 0.01, and 1.00 for models 1, 2, and 3 respectively. In each of these models the true and estimated network correlations are closely related, with few major deviations. The yellow lines show the general relationship using LOESS curves fitted to the points.

### Simulations 2: Statistical power of nodal regression

The results of the nodal regression simulation are shown in Figure 3 for four different network sizes (10, 20, 50, 100). The relationship between network correlation and power is approximately logistic, with a higher power for networks with larger numbers of nodes across the range of network correlations. Large gains in power could be seen for networks with 20 nodes against those with 10 nodes (37% versus 27% respectively, at a network correlation of 50%), whereas networks with 100 nodes against 50 nodes had a much smaller gain in power (55% versus 51% respectively, again at a network correlation of 50%). To attain a statistical power of the conventional 80%, network correlations of 0.81, 0.78, 0.69, and 0.65 were required for network sizes 10, 20, 50, and 100 respectively. The maximum required level of network correlation to achieve 80% statistical power was 0.81 (for *n* = 10).

**Figure 3:**
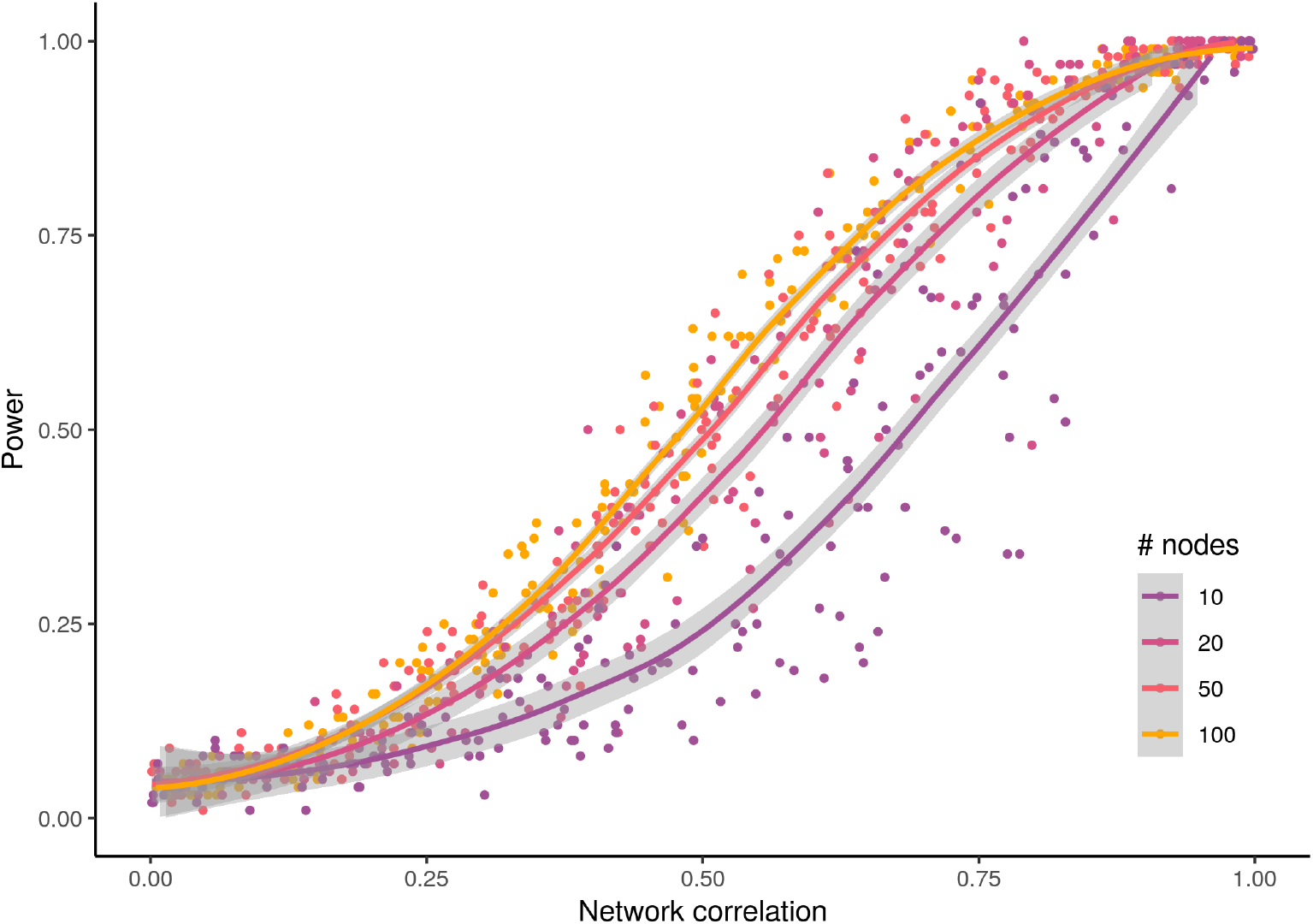
The relationship between estimated network correlation and statistical power for various numbers of nodes over 1000 simulations, where the minimal effect size was chosen such that power was ≥ 99.9%. Power and network correlation are monotonically related in a relationship that resembles a logistic curve. Larger numbers of nodes achieved a higher power for a lower network correlation than smaller numbers of nodes, as is expected in a nodal regression where sample size is equal to the number of nodes. A statistical power of 80% is common by convention, and was generally achieved for all simulated networks with at least 10 nodes for network correlations greater than 0.80.

Table 1 shows the sampling effort *I* required to achieve 80% power, depending on the social differentiation *S* and size of a network *n*. For systems with low levels of social differentiation and low network sizes, high sampling effort (*I* = 530 for *n* = 10, *S* = 0.05) is required. However, as social differentiation increases, much lower sampling effort is required (*I* = 5.3 for *n* = 10, *S* = 0.5). This reflects the findings shown in Figure 1. Larger network sizes also have an effect on the required amount of sampling, with *n* = 100 requiring less than half the sampling effort as *n* = 10.

**Table 1:** Required sampling effort *I* = *µH*(*d*) to achieve a statistical power of 80% for net-works with social differentiation *S* and number of nodes *n*. Relatively low numbers of event observations are required to achieve 80% power for levels of social differentiation of 0.5 and higher.

| $S$ | $n$ | | | |
| --- | --- | --- | --- | --- |
|  | 10 | 20 | 50 | 100 |
| 0.05 | 530 | 410 | 230 | 200 |
| 0.2 | 33 | 26 | 14 | 12 |
| 0.5 | 5.3 | 4.1 | 2.3 | 2 |
| 0.8 | 2.1 | 1.6 | 0.89 | 0.77 |
| 2 | 0.33 | 0.26 | 0.14 | 0.12 |
| 10 | 0.013 | 0.01 | 0.0057 | 0.005 |

### Simulations 3: Optimal network correlation estimator for generic tests

The powers of the nodal regression analyses for this simulation were split into five equally sized groups between 0% and 100%, with the top group of 80 - 100% being the desired power, by convention. The results of the simulations are shown in Figure 4 with the optimal sampling effort overlaid. The optimal sampling effort for low levels of social differentiation (*S* < 0.1) was in excess of a sampling effort of *I >* 500. For slightly larger values of social differentiation (0.1 ≤ *S* ≤ 0.2), the required sampling effort drops quickly from approximately *I* = 500 to *I* = 100. The sampling effort for larger values of social differentiation quickly asymptotes towards zero.

**Figure 4:**
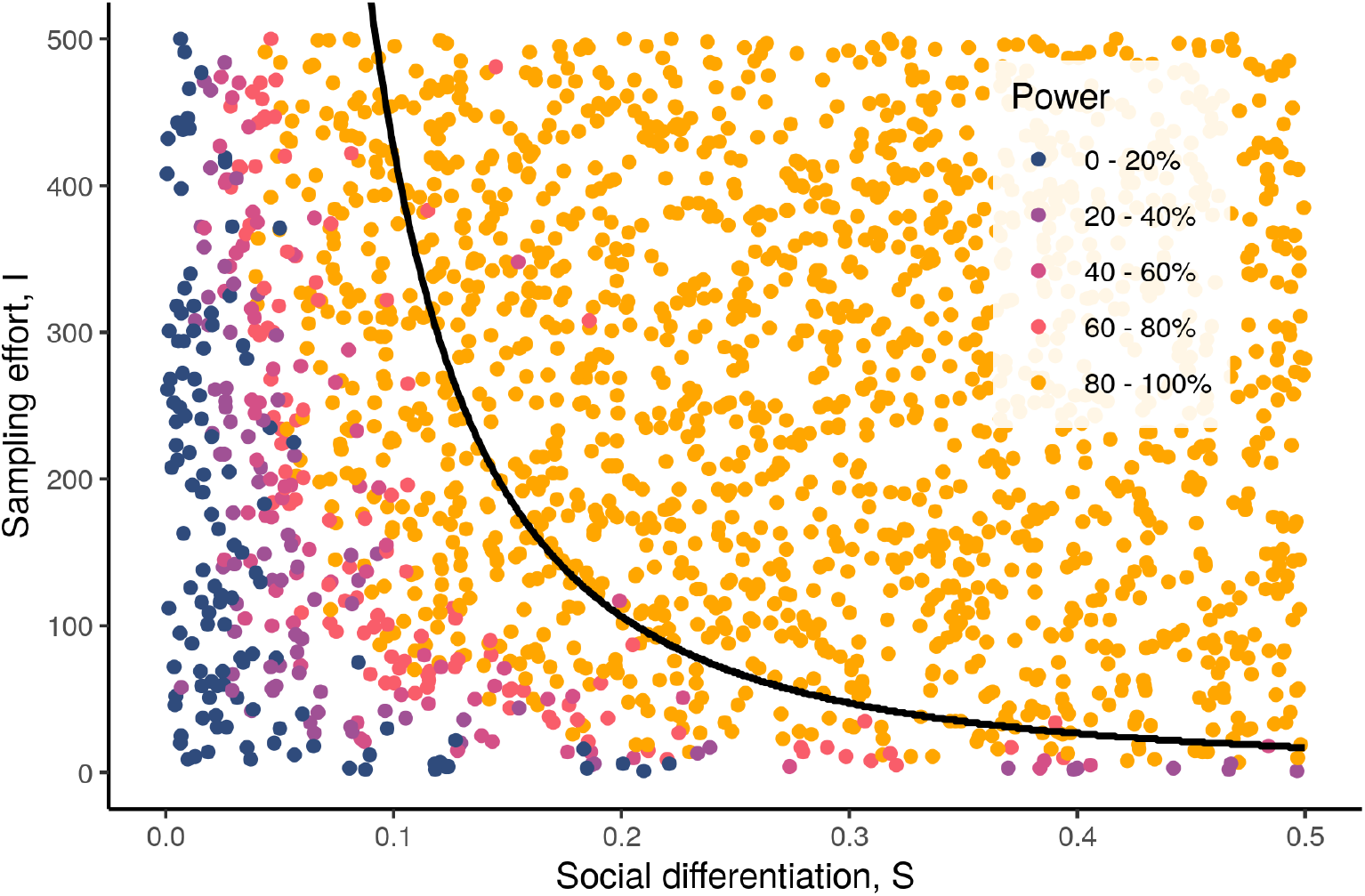
Power of nodal regression analyses for networks with varying levels of social differentiation *S* and sampling effort *I*. The black line shows the optimal sampling effort estimated using the diminishing returns method, corresponding to the optimal network correlation as social differentiation increases. This can be viewed as a classifier where analyses below the line are under-powered, and analyses above the line are adequately- or over-powered.

The optimal sampling effort generally fell around the boundary between power levels of 60 - 80% and 80 - 100% for the nodal regression analyses. Considering the curve as a classifier of under- and over-powered analyses, using a power of 80% as the boundary between the two, 50.3% analyses below the curve were under-powered, and 99.9% of the analyses above the curve were adequately- or over-powered. In the context of our simulated nodal regressions, the curve was relatively conservative, with the power level that most closely corresponded to the power of the optimal network correlations being 90%.

### Case study: Southern resident killer whale contacts

The southern resident killer whale contact network had an estimated network correlation of 0.977%, with a 95% confidence interval of [0.966, 0.985]. Social differentiation was estimated to be 2.54, with a confidence interval of [2.23, 2.89]. According to our power analysis, these levels of social differentiation and network correlation would yield power levels of at least 6.6%, 26.5%, 66.2%, 98.2%, and 100.0% for true effect sizes 0.1, 0.3, 0.5, 0.7, and 0.9 respectively. The diminishing returns estimator suggested that the optimal amount of network correlation to sampling effort is approximately 0.90, corresponding to sampling efforts between 0.51 and 0.86. The true sampling effort was 3.30, several times larger than the maximum recommended sampling effort of 0.86.

## Discussion

In this study we showed that sampling effort can have a major impact on the accuracy of social networks constructed from count data. We showed that this can severely affect the power of statistical analyses, and demonstrated how to carry out power analysis by accounting for sampling effort in both linear regression and generic statistical analysis. We derived an equation to describe how well a sampled network correlates with the true underlying network using a Gamma-Poisson model of event counts. We showed that the equation is a good estimator of the true network correlation, and is robust to the assumptions of the model being moderately broken. We used simulations of nodal regression analyses to find the relationship between network correlation and statistical power. Additionally we developed an estimate for the optimal level of network correlation required for generic analyses by using the concept of diminishing returns, where increases in network correlation come at the cost of increasing levels of sampling effort. We showed that this can be used as an alternative to conventional power analysis by verifying the results against the nodal regression simulations.

The nodal regression simulations suggested that a reasonable level of network correlation to achieve at least 80% power is around 0.80. This value of network correlation has precedent in the behavioural sciences, and would be categorised as “very strong” by Evans (1996). However we arrived at this value through the use of simulations, and would advise careful application of this guideline for different nodal regression-based analyses, and discourage its use on other types of analysis. We believe it is particularly important to note that in our simulations we used the smallest possible effect sizes required to achieve approximately 100% power with infinite sampling. Effect sizes that satisfy this property are highly unlikely in the real world, with the true effect size either being lower, in which case no amount of sampling will be able to obtain full power, or the effect size being higher, in which case lower sampling efforts will be able to obtain full power. For this reason, we suggest using the code we have made available to run a custom power analysis for any specific study.

An alternative method for calculating the required level of network correlation is to use the diminishing returns-based optimal network correlation estimator. In the nodal regression simulations it proved to be a conservative classifier of analysis power, often suggesting that more samples were required than would have been necessary to obtain a power of 80%, and was more in line with a desired power level of 90% in this specific simulation. Since the estimator is based on diminishing returns, it implicitly models the trade-off between lower sampling variance and the cost of sampling, which is a fundamental part of behavioural data collection. For this reason, we believe this generic estimator to be a useful tool for generating estimates of the amount of sampling required, and in making few assumptions, it is flexible, allowing it to be used for any type of social network analysis. Another strength of the method is that knowledge of the desired statistical power is not required. Power can often be difficult to define, and is therefore usually treated as a nuisance parameter when conducting power analyses, with researchers reverting to conventions such as 80%. The free parameter *ρ*_MAX_ could be considered a similar theoretical limitation to this method, but we view it as the maximum value of network correlation a researcher is interested in. We chose *ρ*_MAX_ = 0.99 because the final increase in network correlation by 0.01 from 0.99 to 1.00 is unlikely to be of biological or practical interest, though other values could be used depending on the specific context. We suggest that for nodal regression, the simulations we have provided will be a more precise tool, but for other types of analyses, or where the desired power level is difficult to define, the generic estimator will give a good guide to the amount of sampling needed. The two methods can also be used in conjunction with each other to gain additional understanding about the power of an analysis.

The inspiration for this work, Whitehead (2008), suggested a guideline sampling effort for the social association test of *S*^2^ × *H >* 5, where *H* is the mean number of observed events per individual (Bejder et al., 1998). This is a classifier analogous to the classifier shown in Figure 4 for determining the required level of sampling effort for a given value of social differentiation to achieve a sufficiently-powered analysis. We found that when it came to sampling effort, *I* – Whitehead (2008)’s the number of events observed per dyad – was the important factor for the classifier, whereas (2008)’s guidance suggested that *H* – the number of events per individual – was the important factor. This is a key point because unless a strict sampling regime is used, sampling is unlikely be even across dyads and nodes, and a subset of well-sampled individuals that share connections might be over-sampled, while other dyads remain severely undersampled. This is taken into account by the definition of sampling effort, *I* = *µH*(*d*), since the harmonic mean is lower than the arithmetic mean for unevenly sampled dyads. The previous study showed that this was unimportant in the case of association data on a specific test, but our equation suggests that it should be taken into account for event count data.

We suggest that in addition to conducting power analyses when developing studies or assessing the feasibility of a study, our methods could also be used in post hoc or dynamic social network analyses where subsetting is required. It is common for long-term event data to be split up into multiple networks, often for the purposes of studying changes in social behaviour over time (Hobson et al., 2013). Our methods could be used as a data-driven means to determine the amount of data required for each network, and therefore how many networks should be constructed. In this case a network correlation of 0.99 could be considered to be a representative network by the arguments above, but again this may depend on the context.

Our method uses a Gamma-Poisson model of estimated event rates. This introduces several assumptions that may not be met by empirical data, namely 1) social events occur due to a Poisson point process, 2) true underlying event rates do not change over the course of data collection, and 3) true underlying event rates follow a gamma distribution. Assumptions are necessary for any statistical analysis, and assumptions 1 and 2 are usually assumed implicitly when constructing social networks from count data. Modelling count data as a Poisson point process means that the method will not work with duration data. These assumptions could be broken by sampling biases such as increased observation of gregarious individuals or sampling based on number of social events observed. Assumption 3 is necessary for the method, but may not be met in systems where event rates follow a multimodal distribution. The validity of the assumptions can be tested using the diagnostic tools included in the R package we have provided.

Correlation is a familiar statistical concept, so using network correlation to quantify the impact of sampling on the accuracy of social networks is an attractive method. However, network correlation on its own can only offer limited information about how useful a sampled network will be in further analyses. We used simulations in this study to estimate how network correlation relates to statistical power in nodal regression, but these simulations are limited to specific circumstances, under a number of assumptions. This will always be a limitation of simulation studies, and makes the generality of the results somewhat restricted. We have shown that the required level of network correlation is context dependent. This contrasts with some studies that have used Whitehead (2008)’s method to calculate network correlation, and suggest a minimum acceptable level of network correlation at 0.40 (Chabanne et al., 2017; Hawkins et al., 2020; Frau et al., 2021). Our results suggest a threshold value of 0.40 is not generally optimal when conducting nodal regression on social networks built from count data, and that the true optimal will depend on the level of social differentiation and sampling effort.

We have made the code from this study publicly available, and our method of power analysis can be used for any prospective studies where event count data will be collected, in post hoc analysis when reporting results, to indicate whether a null result may be due to insufficient sampling, and even when subsetting data to ensure sufficient data are used in each network. The diminishing returns method presented here for computing optimal network correlation naturally encodes the cost of collecting behavioural data and provides a simple method for estimating required sampling effort without specifying additional parameters such as desired power. We believe its flexibility could prove it to be useful for a wide variety of social network analyses.

## Author contributions

MNW conceived of the negative binomial network correlation method and JDAH conceived of the power analysis methods. JDAH derived the equations and implemented the simulations with input from MNW, DWF, and LJNB. The manuscript was written by JDAH with input from MNW, DWF, and LJNB.

## Acknowledgments

We would like to thank members of the CRAB Social Network Club and Alexander Mielke for useful discussions on this topic. JDAH acknowledges funding from the Engineering and Physical Sciences Research Council [grant number EP/R513210/1]. LJNB acknowledges funding from a European Research Council Consolidator Grant (FriendOrigins - 864461). DWF and MNW acknowledge funding from the Natural Environment Research Council [grant number NE/S010327/1]. The authors declare no conflict of interest.

## Supplementary Material

### Supplementary material: Derivation of Equation 3

#### Gamma-Poisson model

In Whitehead (2008), the population association indices are assumed to be distributed according to a beta distribution. The beta is defined on [0, 1], making it ideal for modelling association indices that are strictly in the same range. However, when modelling event rates, any positive number of event rates can occur in an interval of time, so we propose using the gamma distribution to model the underlying “true” event rates of the population:

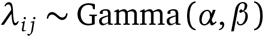

where *λ*_*i j*_ is the event rate between individuals *i* and *j*, and *α, β >* 0 are parameters controlling the shape of the distribution. The observed number of events between *i* and *j, X*_*i j*_ is a an integer count variable, and should therefore be distributed according to a Poisson, depending on the time *d*_*i j*_ spent sampling the dyad (*i, j*):

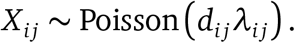

Since *d*_*i j*_ is constant for any particular dya d, the distribution of *d*_*i j*_*λ*_*i j*_ is given by

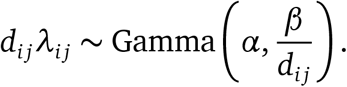

This type of model is known as a Gamma-Poisson mixture, and is equivalent to the conjugate distribution of a Poisson random variable with gamma priors (). The distribution of *X*_*i j*_ can be described by the negative binomial in terms of the gamma parameters *α, β* and the sampling time *d*_*i j*_:

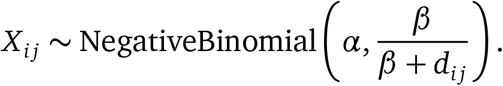

In empirical network studies, the event rate usually used is based on this sampled event count *X*_*i j*_. The estimated event rate 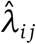 is defined as

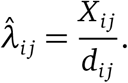

#### Estimating network correlation

The goal of this study is to compute the correlation between the true and estimated event rates *λ*_*i j*_ and 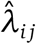 respectively. The Pearson correlation coefficient is defined by

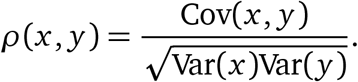

In the same way as Whitehead (2008), we assume that the random variables 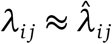, allowing us to set 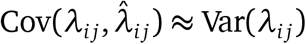, giving the following equation for correlation:

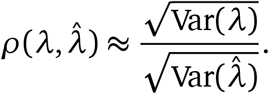

This assumption was verified using the simulations shown in the main study. The variance of the true event rate is simply the variance of the gamma distribution:

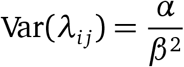

The variance of the estimated event rate is based on the variance of the negative binomial:

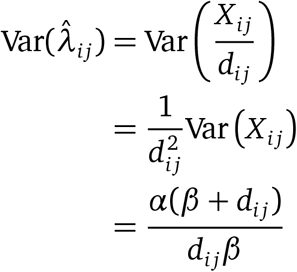

The total variance of the true event rate over the *m* dyads can now be written as:

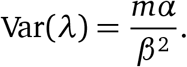

The total variance of the estimated event rates is:

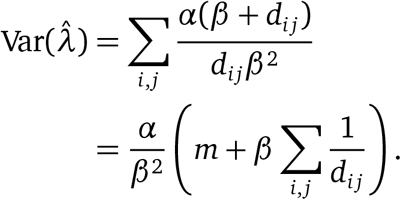

The variance ratio can now be written as:

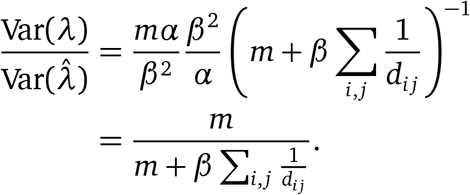

Now the network correlation is given by

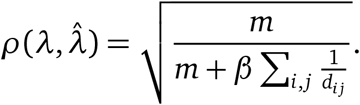

The *α, β* parameterisation of the gamma distribution is not especially biologically meaningful, so we use an alternative parameterisation in terms of the mean event rate *µ* and social differentiation *S*. This parameterisation can be written for *α, β* as:

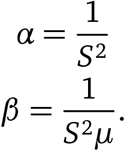

The 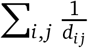 term in the network correlation equation is related to the harmonic mean *H* of the event rates:

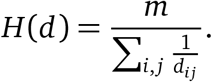

By substituting in these three equations, we arrive at the final equation for network correlation:

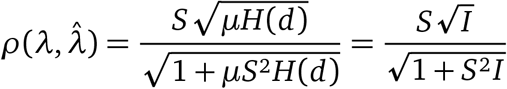

where *I* = *µH*(*d*) is a term introduced to summarise the amount of sampling (we call it the sampling effort) for ease of interpretation.

#### Correlation and accuracy

Correlations aren’t usually a guarantee of accuracy. For example, the correlation between *x* and 2*x* is 1.0, but *x* is mostly a very inaccurate estimate of 2*x*. However in our context, the correlation 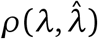 *can* be interpreted as a measure of accuracy. To demonstrate this fact, consider a common measure of accuracy, the mean squared error:

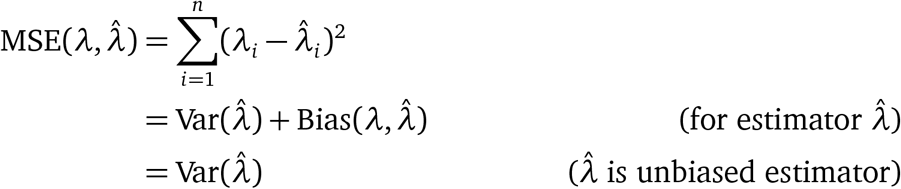

Now consider the correlation from earlier, the square root of the variance ratio:

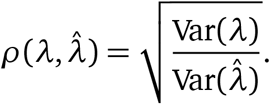

Since Var(*λ*) is fixed, this means that as 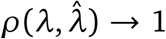, then 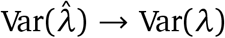; and conversely that as 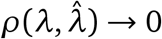, then 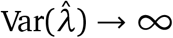. Therefore, increases in correlation reduce the mean squared error, and increase accuracy. On the other hand, decreases in correlation increase the mean squared, and decrease accuracy.

An equivalent measure of accuracy is the mean absolute error, which is the square root of mean squared error. This definition gives:

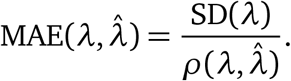

Since SD(*λ*) is fixed for any given network, then the mean absolute error and network correlation are inversely related.

#### Likelihood estimator

To estimate the parameters of the gamma distribution, we cannot simply calculate the mean event rate and social differentiation of the estimated event rates, since they are subject to sampling error. Therefore we use our Gamma-Poisson model to estimate the parameters of the underlying “true” distribution of event rates. Unfortunately the maximum likelihood estimator of the negative binomial only exists for data where the sample variance is larger than the sample mean. This could be limiting in our case, so instead we use numerical maximum likelihood estimation. The log-likelihood of our model is derived from the likelihood of the negative binomial, and is given by

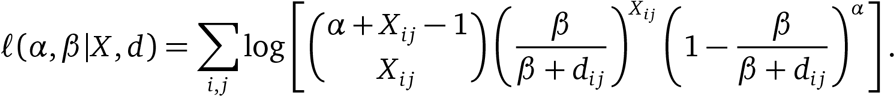

We used a numerical method (Nelder-Mead) to maximise the log-likelihood and estimate the parameters *α, β*. The parameters *µ, S* were calculated from these. Alternative methods such as normal approximation or Gibbs sampling can also be used to estimate the parameters, and would provide confidence*/*credible interval estimates for the network correlation. We did not use these methods in our simulations because of the potential for estimation errors, but have included them in the package and code, and recommend their use for typical, one-off analyses where the diagnostics can be checked.

#### Normal approximation of confidence intervals

To estimate the confidence intervals of *α* and *β* we used a normal approximation. The normal approximation assumes the parameters are distributed according to a multivariate normal distribution. By the Bernstein-von Mises theorem, we can approximate the confidence interval by drawing samples from a multivariate normal distribution:

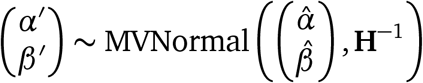

where *α’, β’* are samples from a multivariate normal distribution centered at the maximum likelihood parameter estimates 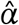 and 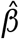 with covariance equal to the inverse Hessian matrix **H**.

Confidence intervals can be estimated from this distribution by sampling *α*^’^ and *β*^’^ repeatedly and calculating percentiles for each parameter. The R code to do this is shown below:

~~~
# Likelihood function for negative binomial model.
lk_nbinom <- function(par, x, d) {
 par <- exp(par)
 r <- par[1]
 p <- par[2]/(par[2] + d)
 sum(dnbinom(x, r, p, log=TRUE))
}
# Estimate parameters using optim from data x and d.
optim_obj <- optim(
 c(0, 0),
 function(par) -lk_nbinom(par, x, d),
 hessian=TRUE
)
# Generate samples of parameters using quadratic approximation.
parameters <- exp(
mvrnorm(1e5, optim_obj$par, solve(optim_obj$hessian))
)
a_ <- parameters[,1]
b_ <- parameters[,2]
# Compute social differentiation and event rate.
S <- 1/sqrt(a_)
mu <- a_/b_
# Compute sampling effort and network correlation.
I <- mu * .hmean(d)
rho <- (S * sqrt(I))/sqrt(1 + S^2 * I)
# Quantiles of sampling effort and network correlation.
S_q <- quantile(S, probs=c(ci_lower, 0.5, ci_upper))
rho_q <- quantile(rho, probs=c(ci_lower, 0.5, ci_upper))
mu_q <- quantile(mu, probs=c(ci_lower, 0.5, ci_upper))
~~~

### Supplementary material: Details of simulations

Simulations were carried out by sampling counts of social events from generative models that follow or break the assumptions of the methods to varying degrees. By sampling from a generative model, we are able to control the true event rate (SRI) precisely.

Our simulations assume that there are no sampling biases affecting the probability of seeing some events relative to others. Though this is unlikely to be true for many empirical systems, the SRI makes the same assumption. In the presence of sampling biases, the network itself will be affected by the same biases, and without knowledge of how the biases arise, it is challenging to correct for them statistically. Several approaches have been developed that attempt to correct for this in various types of analysis, but are outside the scope of this paper ().

#### Simulations: Scenario 1

This simulation is designed to follow the assumptions of the model exactly. Social events between pairs of individuals are assumed to be drawn from a Poisson process with a fixed rate of occurrence. The underlying rates are modelled according to a Gamma distribution parameterised by social differentiation *S* and *µ*. Sampling times *d*_*i j*_ were uneven, and were drawn from a Poisson distribution with rate *t* ∼ *U* (1, 10). Any sampling times *d*_*i j*_ equal to 0 were set to 1, to avoid division-by-zero errors.

The full generative model is given below:

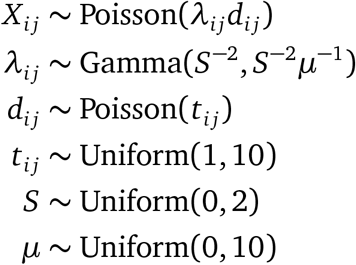

#### Simulations: Scenario 2

This simulation is designed to simulate the effect of community structure on the method. Community structure could be a problem because the underlying distribution may sometimes be multi-modal, and therefore not modelled well by the Gamma distribution. For simplicity we did not simulate nodal membership of different groups, and instead focused on the issue of a bimodal underlying distribution. This does not limit the generalisability of the simulation because the method does not model dyads according to their nodal attachments, and instead models each dyad as a draw from a Gamma distribution. To simulate a multi-modal distribution, we categorised dyads into two classes: class (1), between members of the same community; and class (0), between members of different communities. The probability of being assigned to class (1) is given by *π* ∼ *U* (0, 1). Dyads in class (1) have an event rate *µ*_1_ ∼ *U* (0, 10), and dyads in class (2) have a lower event rate *µ*_0_ ∼ *U* (0, *µ*_1_).

The full generative process is given below:

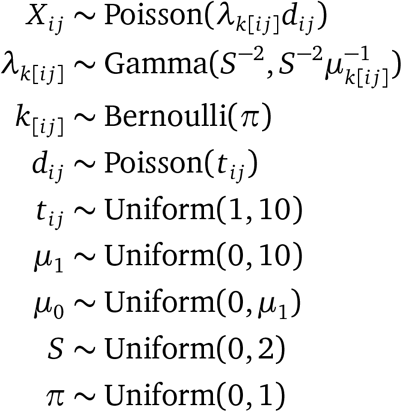

#### Simulations: Scenario 3

This simulation studies the effect of zero-inflated event rates, where a dyad’s SRI can be zero due to either insufficient sampling or because the underlying social preference is zero. All other features of the original scenario are kept the same, but event counts are modelled as coming from a zero-inflated Poisson distribution with zero-inflated parameter *π*.

The full generative model is given below:

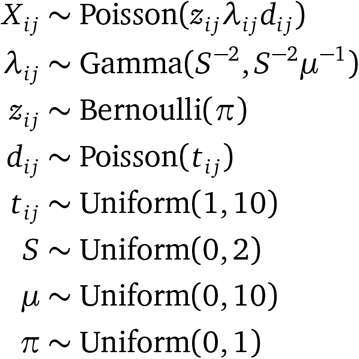

### Supplementary material: Additional verification simulations

The model verification simulations were run for additional scenarios not shown in the main text.

#### Beta Poisson

The gamma distribution is a natural choice for modelling lower-bounded rates thanks to its flexibility, however, it is technically possible for rates to be limited by an upper-bound. To test how well Equation 3 estimates network correlation in this case, we used a Poisson model of event counts with event rates drawn from a beta distribution with a given mean *µ* and social differentiation *S*.

**Figure 1:**
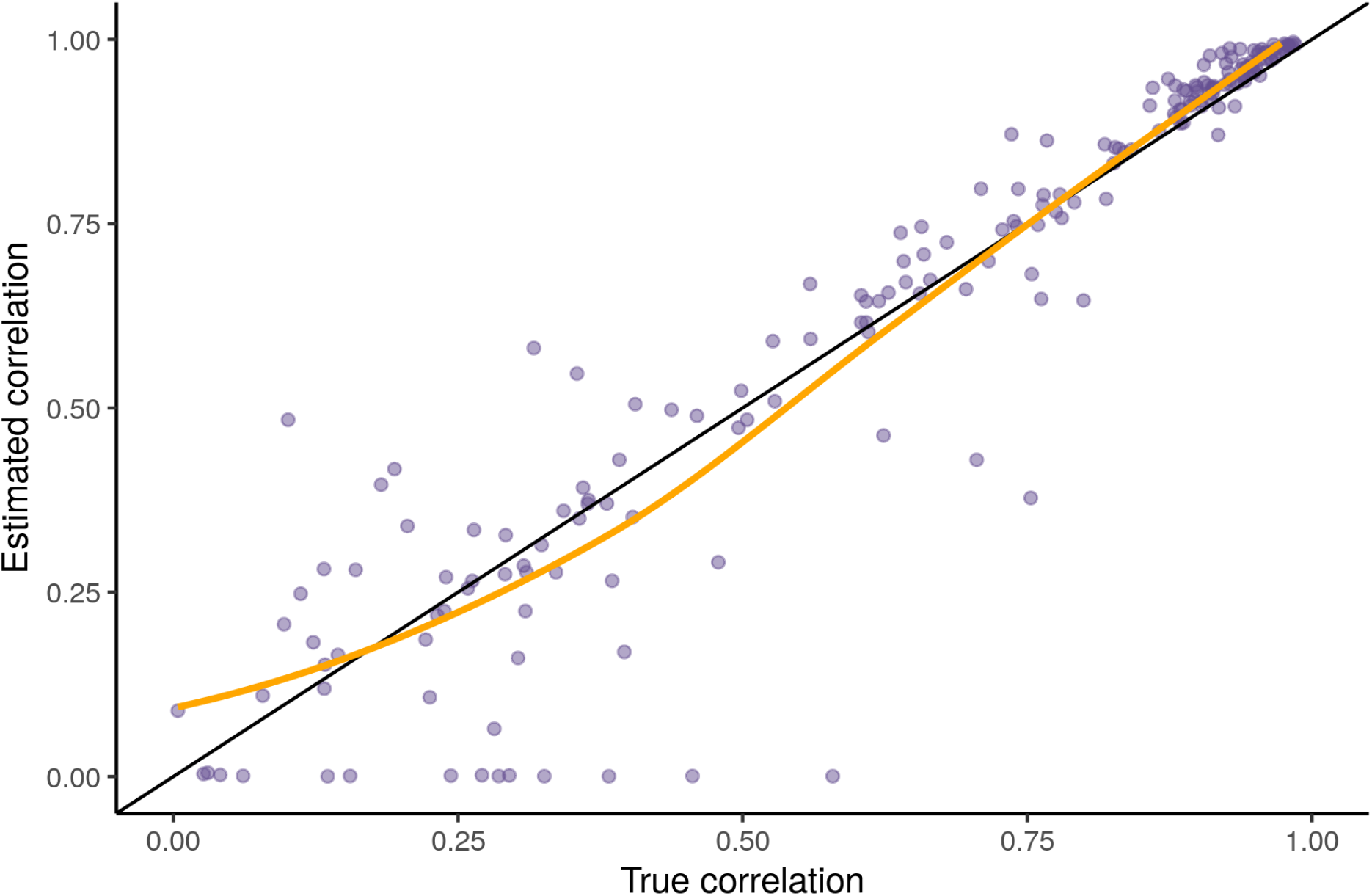
Estimated network correlation against true network correlation in the zero-inflated Poisson scenario. The estimated network correlation values had a correlation of 96% with the true correlation values, and a mean absolute error of 3.5%.

Equation 3 performed well in the beta Poisson scenario. This might be because the gamma distribution can be similar in shape to the beta. This simulation confirms that this method of estimating network correlation will work reasonably well for upper-bounded rates.

#### Beta binomial

The original paper (Whitehead, 2008) assumes the data are distributed according to a binomial distribution with parameters given by a beta distribution. This is often relevant when using association indices such as the simple ratio index to estimate the proportion of time two individuals associate. With this type of data we recommend using the original method. However, we implemented a simulation of this kind of data to test how well our network correlation estimate fares when its assumptions are severely broken. In these simulations we set the mean event rate *µ* and social differentiation *S* in the same way as the previous simulations. The results are shown in Figure 2.

**Figure 2:**
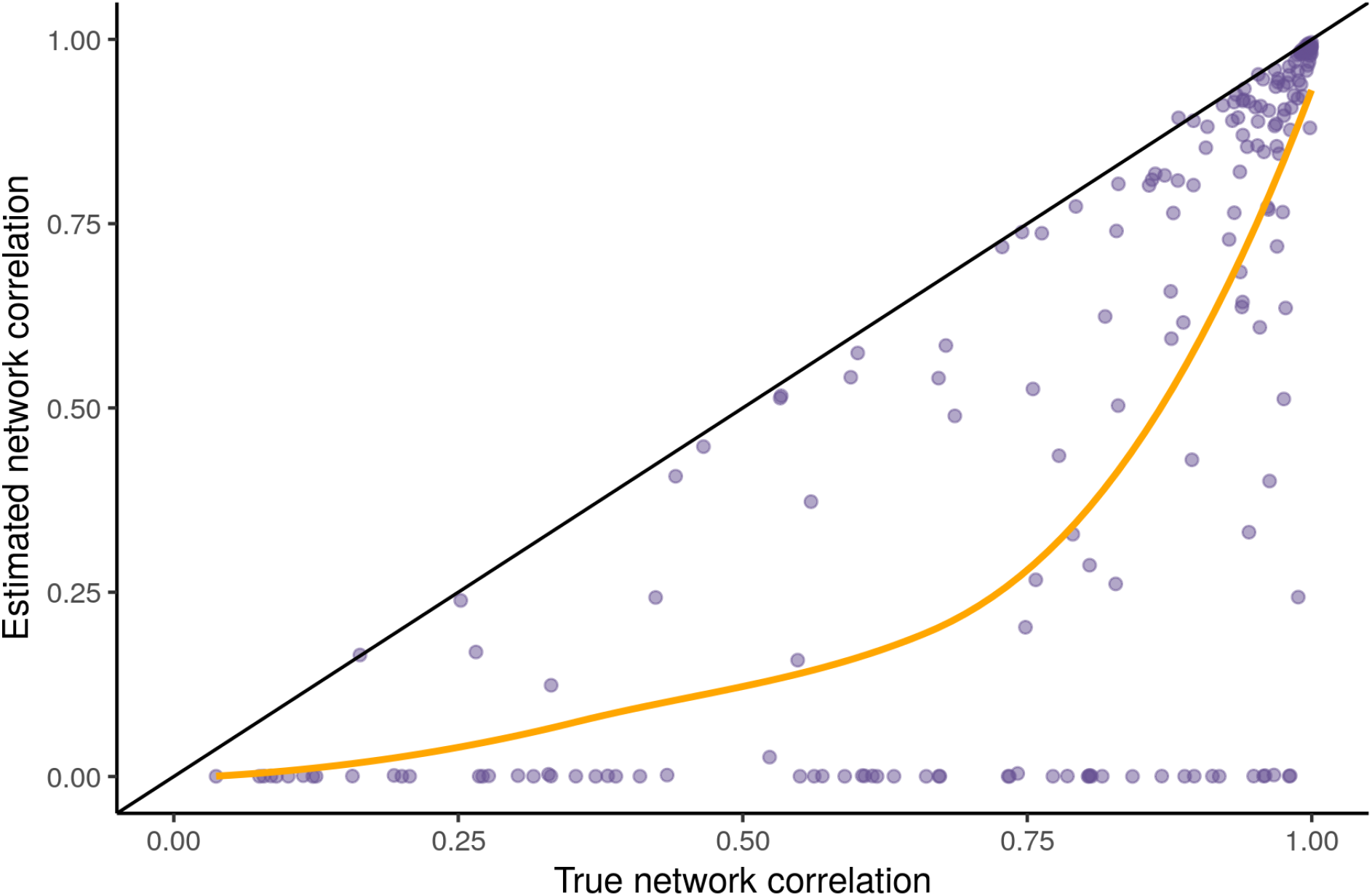
Estimated network correlation against true network correlation in the Beta binomial scenario. The network correlation estimator performs poorly in this scenario, consistently underestimating the true network correlation. In many cases a network correlation of *ρ* = 0 was predicted where the true network correlation was anywhere between 0 and 1.

The poor performance of Equation 3 in this scenario was expected, since the simulation generates data consistent with binary sampling, whereas our network correlation estimate assumes event count sampling. This means that most of the underlying assumptions are broken, making this equation for network correlation not applicable to this type of data. This confirms that when using binary sampled data, Whitehead (2008)’s method should be used to calculate network correlation, but when using event rate data, our Equation 3 should be used.

#### Uneven sampling

In many cases, sampling is uneven across individuals, and consequently some dyads may be more heavily sampled than others. Any unevenness in sampling should not affect our method since the harmonic mean, which is used to determine sampling effort (*I* = *µH*(*d*)), is sensitive to imbalanced values, and will naturally capture uneven sampling. To check this, we created a simulation like scenario 1, but where the number of samples *d*_*i j*_ is drawn from one of two Poisson distributions, one with mean *D*_low_ ∈ [1, 10], and another with mean *D*_high_ ∈ [10, 100].

**Figure 3:**
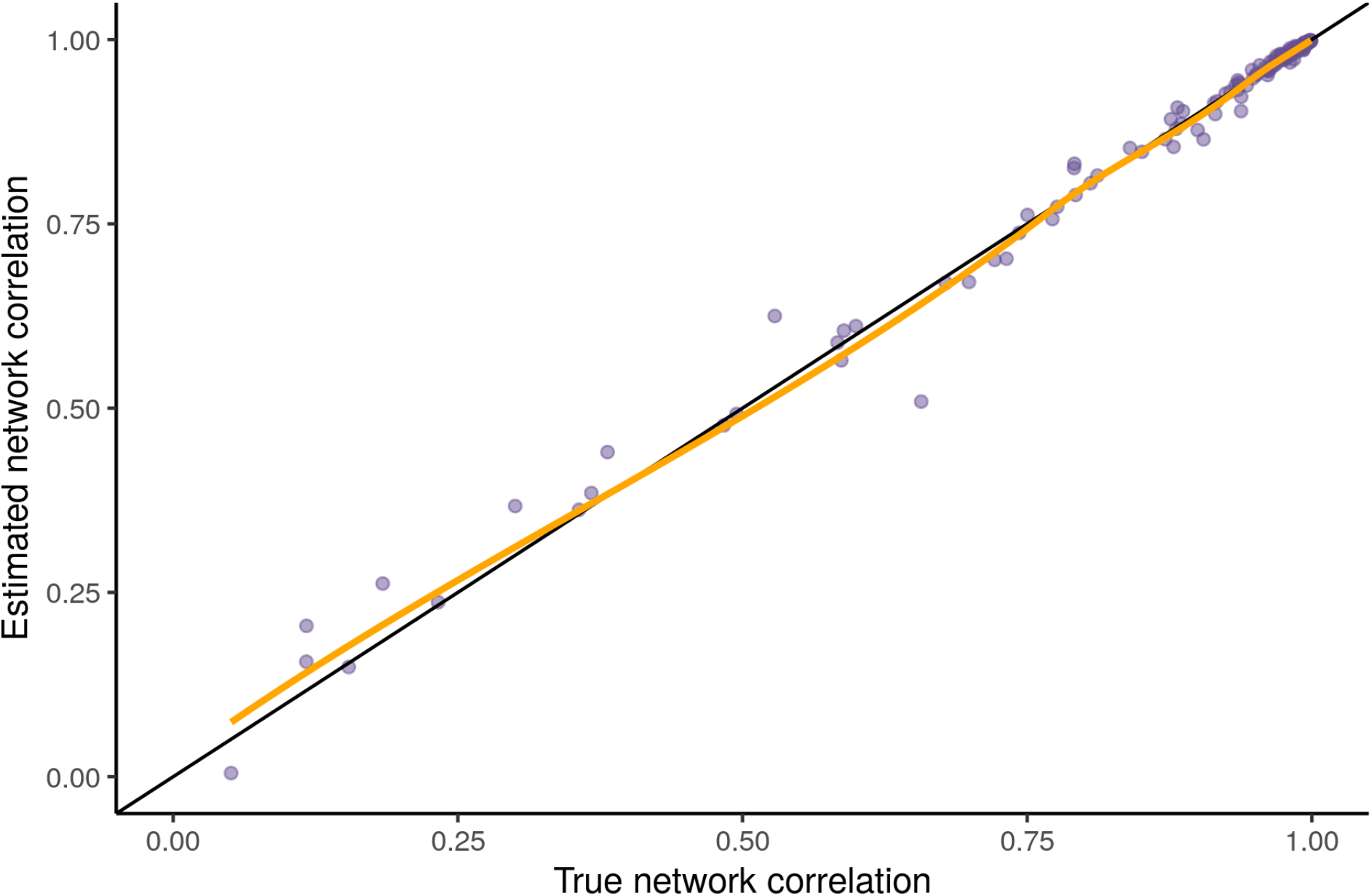
Estimated network correlation against true network correlation in the uneven sampling scenario. The network correlation estimator performs well in this scenario since none of the assumptions of the model are being broken.

The estimator performed well in this scenario, with a mean absolute error of 0.7% and an overall network correlation of 99.5%. None of the assumptions of the model were broken by this scenario, so the estimator was expected to perform well here. This demonstrates that uneven sampling will not affect the network correlation estimate.

### Supplementary material: Elbow computation algorithm

To find the elbow point of diminishing returns, we rotate the curve of *ρ* against *I* by *θ* radians about the origin (0.0, 0.0), and find the maximum point of the rotated curve. The angle *θ* would ideally be determined by the maximum point of the curve, but in our case, *ρ* is asymptotic to 1.0, so instead we introduce a free parameter *ρ*_MAX_ that describes the effective maximum correlation.

Mathematically, to rotate a point (*x, y*) *θ* degrees about (0, 0) we can use the rotation matrix to get the new points (*x’, y’*) in the following way:

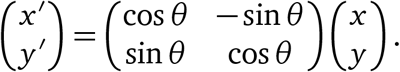

By doing this for all points of the curve, we get a rotated version of the original curve. The maximum point of the transformed curve is the elbow point of the curve.

~~~
I <- seq(0, 100000, 1) # Possible values of sampling effort I.
# Estimated social differentiation, set to 2.5 for example.
S <- 2.5
# Calculate correlation for values of I.
rho_ <- rho(S, I)
# Find effective max of curve using threshold.
rho_argmax <- which.min(abs(rho_ - threshold))
max_rho_ <- rho_[rho_argmax]
max_I <- I[rho_argmax]
# Calculate rotation matrix to rotate curve around (0, 0).
theta <- atan2(max_rho_, max_I)
co = cos(theta) si = sin(theta)
rotation_matrix = matrix(c(co, -si, si, co), nrow=2, ncol=2)
# Rotate curve around (0, 0).
rotated_data <- cbind(rho_, I) %*% rotation_matrix
# Optimal rho, by diminishing returns principle.
rho_prime <- rho_[which.max(rotated_data[, 1])]
# Corresponding optimal sampling effort.
I_prime <- I[which.max(rotated_data[, 1])]
~~~

### Supplementary material: Nodal regression with eigenvector centrality

In the main text we used strength to assess the power of nodal regression, but the choice of network metric should not affect the results since the effect size was calculated relative to the metric when estimating the effect size required for ≈ 100% power. To verify this, we ran the simulations with eigenvector centrality as the network metric. These simulations replicated our results, showing that the results should hold for any custom power analysis run using our code.

**Figure 4:**
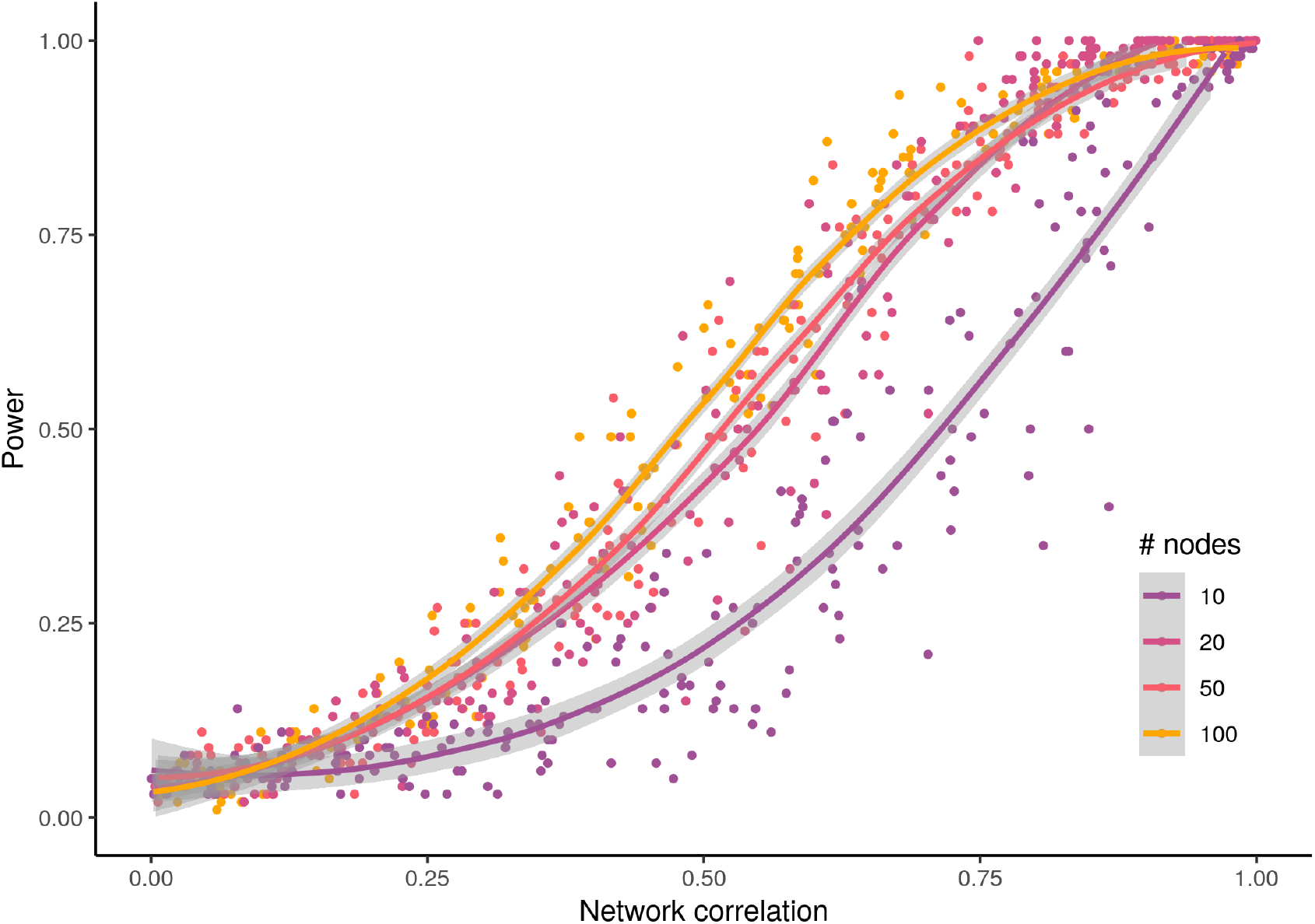
Relationship between power and network correlation for different numbers of nodes in nodal regression simulations with eigenvector as the predictor variable.

### Supplementary material: Effect of *ρ*_MAX_ on optimal network correlation estimate

The generic diminishing returns method relies on a free parameter *ρ*_MAX_ indicating the effective maximum amount of network correlation that is biologically relevant. In the main text we used *ρ*_MAX_ = 0.99, but any value between 0 and 1 could theoretically be used. Values of *ρ*_MAX_ close to 0 might not be advisable, but there is scope for parameter choice for values of *ρ*_MAX_ closer to 1. To evaluate the effect of parameter choice on the classifier, we repeated Simulation 3 for three value: *ρ*_MAX_ ∈ {0.9, 0.99, 0.999}. These values were chosen to have different orders of magnitude difference to 1 to show the realistic maximum range of the classifier.

**Figure 5:**
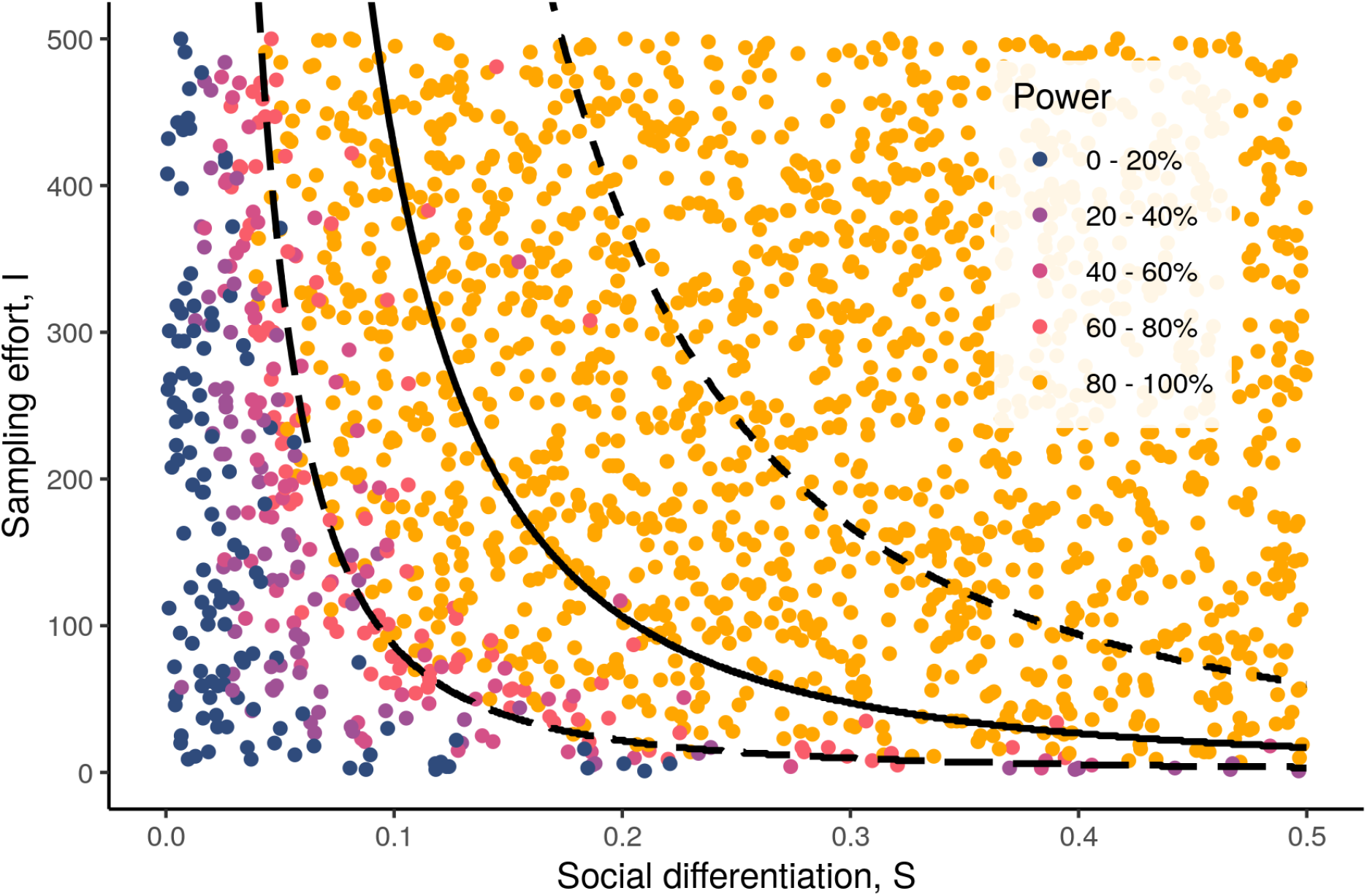
Simulation 3 was re-run for three different values of *ρ*_MAX_ ∈ {0.9, 0.99, 0.999} to test the sensitivity of the diminishing returns method to parameter choices with different orders of magnitude difference to 100% network correlation. The solid black line indicates the value of *ρ*_MAX_ = 0.99 used in the main text, the long dashed line indicates *ρ*_MAX_ = 0.9, and the short dashed line indicates *ρ*_MAX_ = 0.999.

The parameter choice makes of *ρ*_MAX_ a significant impact on the classifier. A parameter value of *ρ*_MAX_ = 0.90 yielded a classifier closely matching the 80% power boundary for the nodal regression simulations, whereas the higher value *ρ*_MAX_ = 0.999 was extremely conservative, suggesting much higher sampling efforts than required to attain a high level of power. These results are difficult to extrapolate to other contexts, but by keeping the definition of *ρ*_MAX_ as the maximum network correlation of biological relevance, this method can be used to provide guidance on the required sampling effort for a given level of social differentiation.

### Supplementary material: Case study code

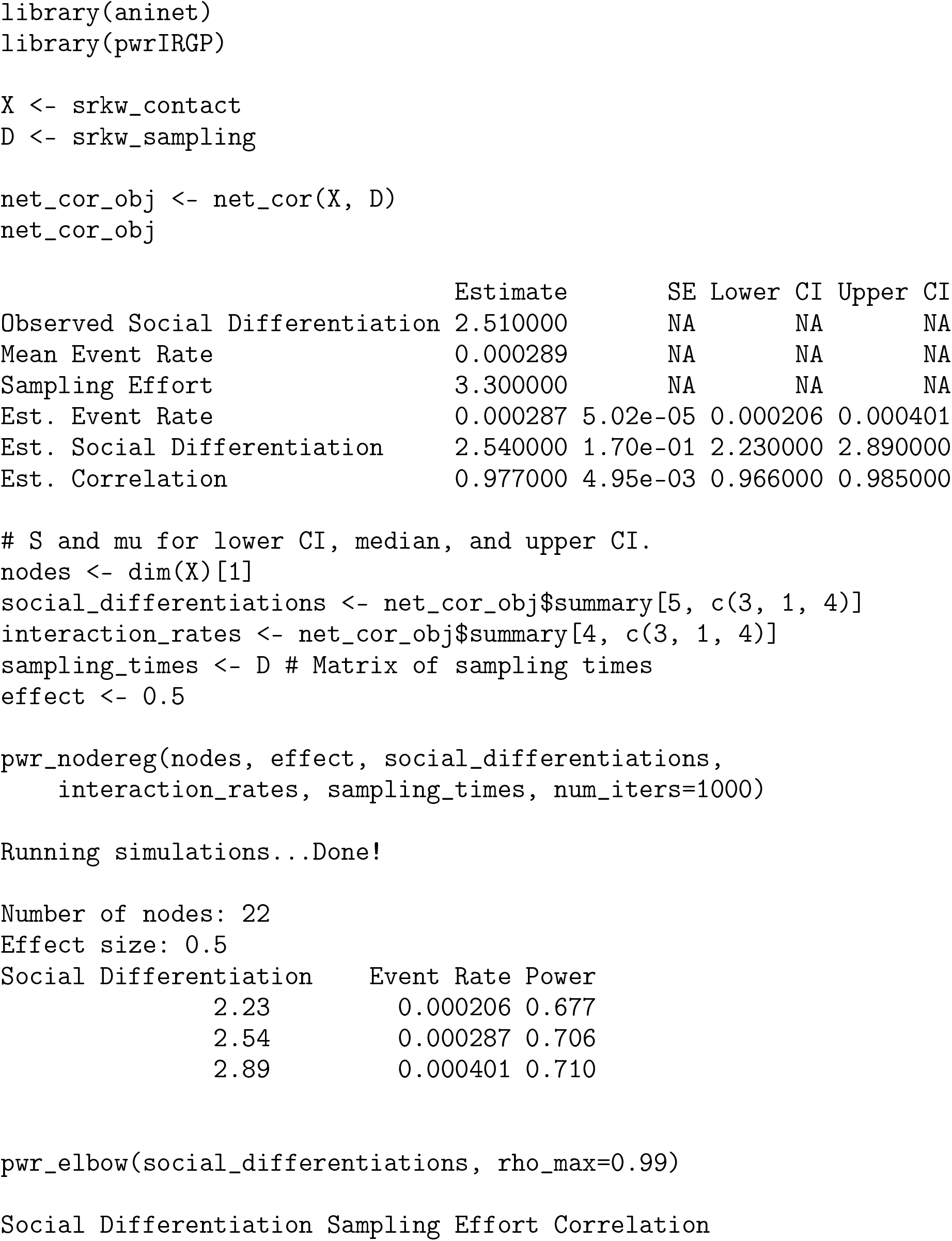
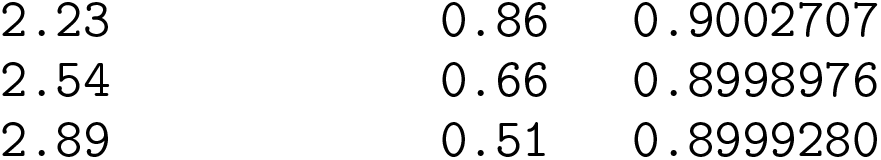

